# MODA: MOdule Differential Analysis for weighted gene co-expression network

**DOI:** 10.1101/053496

**Authors:** Dong Li, James B. Brown, Luisa Orsini, Zhisong Pan, Guyu Hu, Shan He

## Abstract

Gene co-expression network differential analysis is designed to help biologists understand gene expression patterns under different conditions. We have implemented an R package called MODA (Module Differential Analysis) for gene co-expression network differential analysis. Based on transcriptomic data, MODA can be used to estimate and construct condition-specific gene co-expression networks, and identify differentially expressed subnetworks as conserved or condition specific modules which are potentially associated with relevant biological processes. The usefulness of the method is also demonstrated by synthetic data as well as Daphnia magna gene expression data under different environmental stresses.

## 2 INTRODUCTION

Gene co-expression network attracts much attention nowadays. In such a network, nodes represent genes and each edge connecting two genes stands for how much degree may this pair of genes are co-expressed across several samples. The presence of these edges is commonly based on the correlation coefficients between each gene pairs. The higher of correlation between a pair of genes, the higher probability that there exists a co-functionality relationship between them. Weighted correlation network analysis (WGCNA) [1, 2] has been widely used for this case, mostly as a tool for single network analysis. Traditional gene differential analysis has covered identification of important individual genes [3] which shows significant changes across multiple conditions. Even so-called network differential analysis still focus on isolated nodes (genes) in networks. However, based on the fact that genes interact with each other to exert some biological function instead of acting alone, it may be more informative to identify a subnetwork of genes which are conserved across multiple conditions or just active in certain conditions.

Several previous works went beyond individual gene differential analysis. GSCA[4] detects a set of differentially co-expressed (DC) genes. DICER [5] uses a probabilistic framework to detect DC gene sets. Both of take genes as individuals and did not provide a systematic view (at network level) of expression profiles. DINGO[6] estimates group-specific networks by calculating differential scores between each pair of two genes, which is focused on individual edges in the networks.

Here we present MODA (Module Differential Analysis), a Bioconductor (ref Huber) package for co-expression network differential analysis, which can (i) estimate and construct condition-specific co-expression networks with limited samples for each biological condition from gene expression profile; (ii) identify conserved and condition-specific co-expression modules by comparing networks; (iii) perform functional annotation enrichment analysis on the identified modules. We first validated our MODA on synthetic data set and then applied it to *Daphnia magna* gene expression data under different environmental stresses.

## 3 Methods

The first step is condition-specific network reconstruction from gene expression profile. In a gene co-expression network, the edge weights are defined by correlation coefficients of gene pairs. However, it is well known that the accurate correlation coefficient is approximated by 1/*sqrt*(*n*) where *n* is the number of samples, which makes it impossible to get reliable correlation coefficients with only several replicates under each experimental condition in practice. We use a sample-saving approach to construct condition-specific co-expression networks for each single condition, which works as follows. Assume network *N*_1_ is background, normally containing samples from all conditions, is constructed based on the correlation matrix from all samples. Then condition *D* specific network *N*_2_ is constructed from all samples minus samples belong to certain condition *D* [7]. The differences between network *N*_1_ and *N*_1_ is supposed to reveal the effects of condition *D*. The rationale behind this criteria is based on the mechanism of correlation, i.e. which samples can make impact on the correlation coefficient while others may not? More details can be found in supplementary file part 1. Finally we get a set of condition specific networks as such.

The second step is module identification for each network. Similar to WGCNA, we also employ hierarchical clustering as the basic method [1, 2]. However, in order to obtain good module identification results, it is crucial to set an optimal cutting height of hierarchical clustering tree, which is usually tune by the users in WGCNA. In our MODA, we propose an automatic method to determine the optimal cutting height based on the quality of modules. Inspired by the concept of partition density of link communities [8], our method search for the optimal cutting height which maximize the average density of resulting modules. Here we simply define the module density as the average edge weights in one module (Equation 1 in supplementary file), same as in [2]. We also provide other criterion such as average modularity for weighted network [9] of resulting clusters to determine the cutting height.

The third step is network differential analysis. We compare two networks by comparing two set of modules. The similarity of each pair of modules is measured by a Jaccard similarity coefficient. With all condition-specific networks compared with the background network, we get a similarity matrix *A*, where each entry *A_ij_* means the Jaccard similarity coefficient between the *i*-th module from the network *N*_1_ and *j*-th module from the network *N*_2_. Then the elements in row sum of *A* (vector denoted by s) indicate how much degree that modules in *N*_1_ can be affected by corresponding condition. The higher **s**_*i*_ means the module *i* in *N*_1_ keeps relatively high row sum of *A* compared with all other *N*_2_ (remove one condition each time), showing these modules have little association with any specific conditions. While lower **s***i* means module *i* in *N*_1_ is very different from the modules in *N*_2_, which may indicate the module has some connection with corresponding condition.

After determining which module may be condition specific or conserved, our MODA package also provides the functionality to associate biological processes with the identified conserved and condition-specific modules by functional annotation enrichment analysis. This is achieved by integrating DAVID [10] based on an R Webservice interface [11]. Figure 1 shows the general process of each step mentioned above.

**Figure 1:**
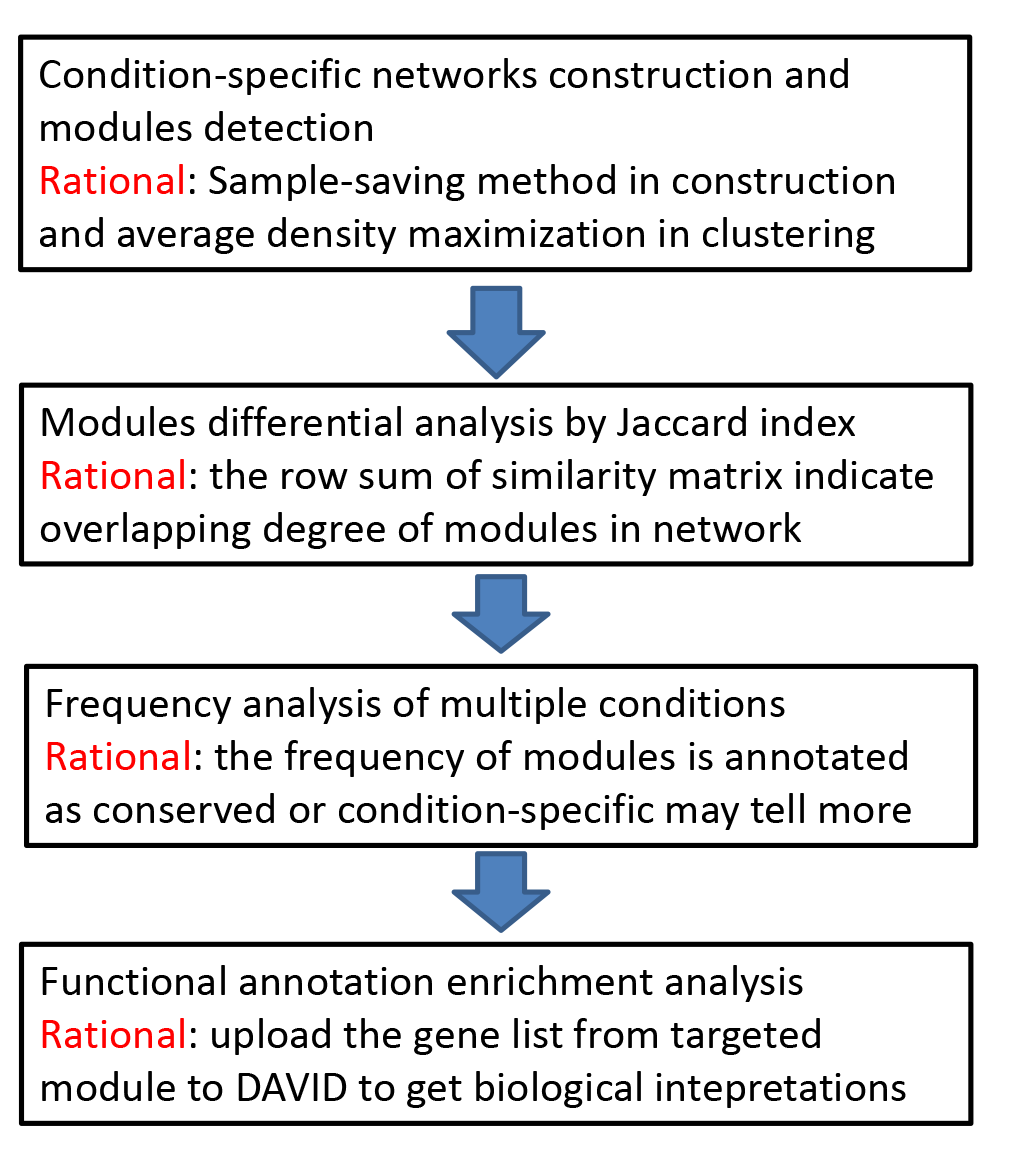
Overview of MODA.

## 4 Result

We evaluated the effectiveness of proposed methods on both synthetic data and real-world data. By comparing two gene expression profiles generated by different desired correlation matrices of the same set of genes, we can determine the genes affected by a groups definition, which is consistent with the generator. The details for simulation as well as the usage of package can be found in supplementary file part 2. The method is also used on a comprehensive RNA-Seq data set obtained from two natural genotypes fo D. magna, to detect condition-specific as well as conserved responsive genes and biological functions. Several biological meaningful results show the capability of this package, and more details can be found in *Characterization of early stress transcriptional response in the waterflea Daphnia magna* (in preparation).

## 5 Supplementary

### 5.1 Concept part

Given gene expression profile *X* ∊ ℝ*^n×p^*, where *n* is the number of experimental samples and *p* is the number of genes. *X_ij_* means the expression value of the *j*-th gene in *i*-th sample. The popular tool WGCNA [1] conducts the module detection by hierarchical clustering, i.e. putting similar gene together. The definition of similarity ranges from basic correlation to more complex topological overlap measure [2]. While how to determine the cutting height of hierarchical clustering tree remains an open problem. Here we give the option to chose the height based on the quality of partition. Inspired by the concept of partition density of link communities [8, 12], we choose the cutting height to make the average density of resulting modules to be optimal. The density of one module *A* is defined as:
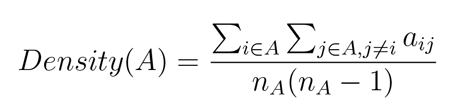
where *a_ij_* is the similarity between gene *i* and gene *j*, and *n_A_* is the number of genes in *A*. We can also use the modularity *Q* of weighted network *A* [9] as the criterion to pick the height of hierarchical clustering tree:

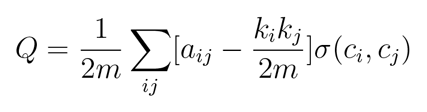
where *m* is the number of edges and *k_i_* is the connectivity (degree) of gene *i*, defined as Σ*_j_a_ij_*. And σ(*c_i_*, *c_j_*) = 1 only when gene *i* and *j* are in the same module. The complete module detection and average density is shown in Figure 2.

After the module detection, the co-expression network is represented as a collection of modules (see Figure 3), which makes the differential analysis more focused on the modules other than the nodes or links. By comparing all module pairs from *N*_1_ and *N*_2_, we can get a similarity matrix *B*, where each entry *B_ij_* means the similarity between the *i*-th module from the network *N*_1_ (denoted by *N*_1_(*A*_*i*_)) and *j*-th module from the network *N*_2_ (denoted by *N*_2_(*A_j_*). The similarity is evaluated by the Jaccard index.

**Figure 2:**
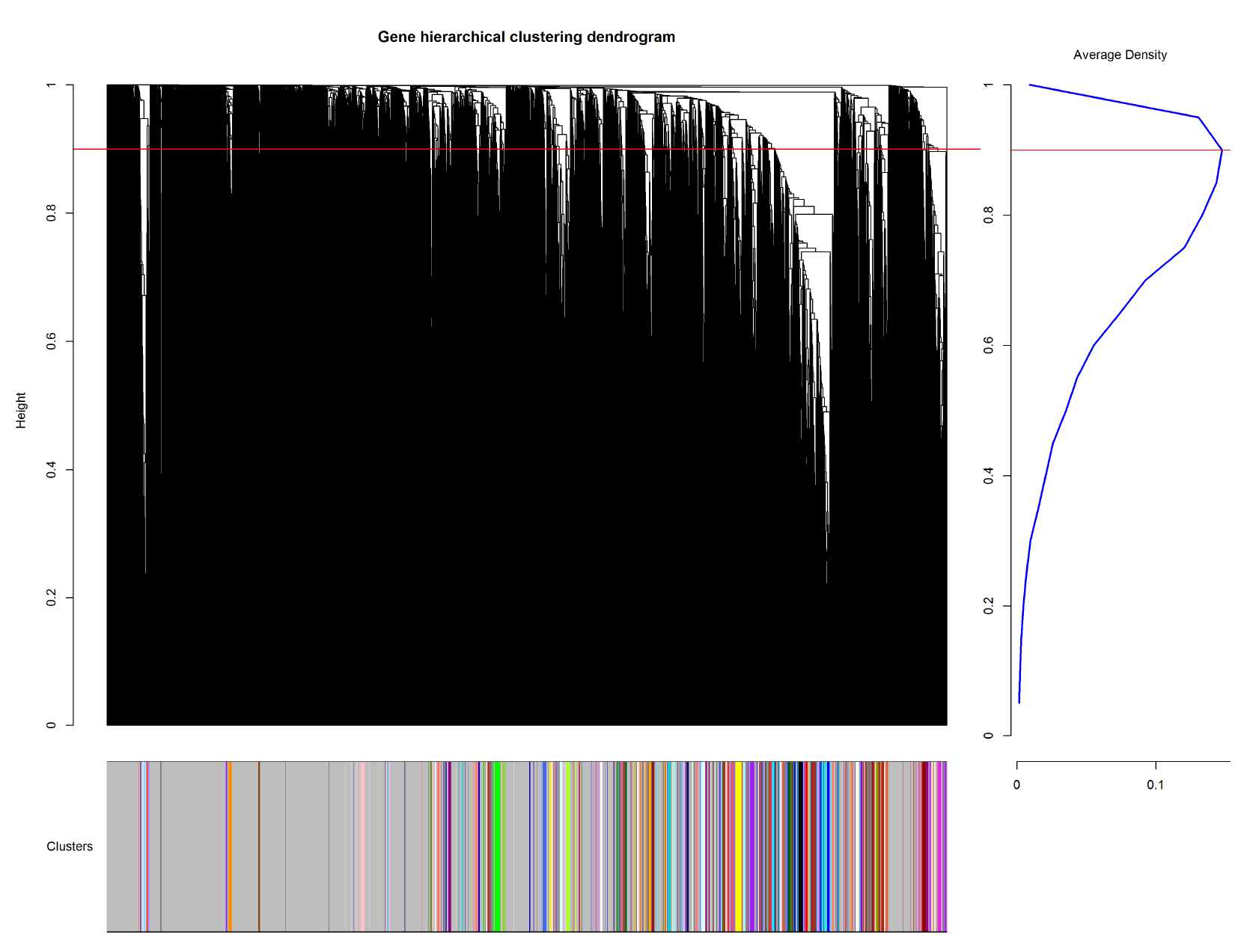
Maximal partition density based hierarchical clustering

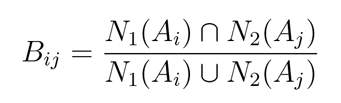

Assume *N*_1_ is background, normally containing samples from all conditions, and the *N*_2_ is constructed from all samples except samples belonging to certain condition *D*. Let **s** is the sums of rows in *B*, i.e. **S**_*i*_ = Σ_*j*_*B*_*ij*_. The value of **S**_*i*_ indicates how much the *i*-th module from network *N*_1_ might be affected by condition *D*. The rationale behind this statistics is based on the mechanism of correlation, i.e. which samples could make an impact on the correlation while others may not? Figure 3 illustrates an extreme example about how the additional two samples may affect the correlation between *X* and *Y*.

**Figure 3:**
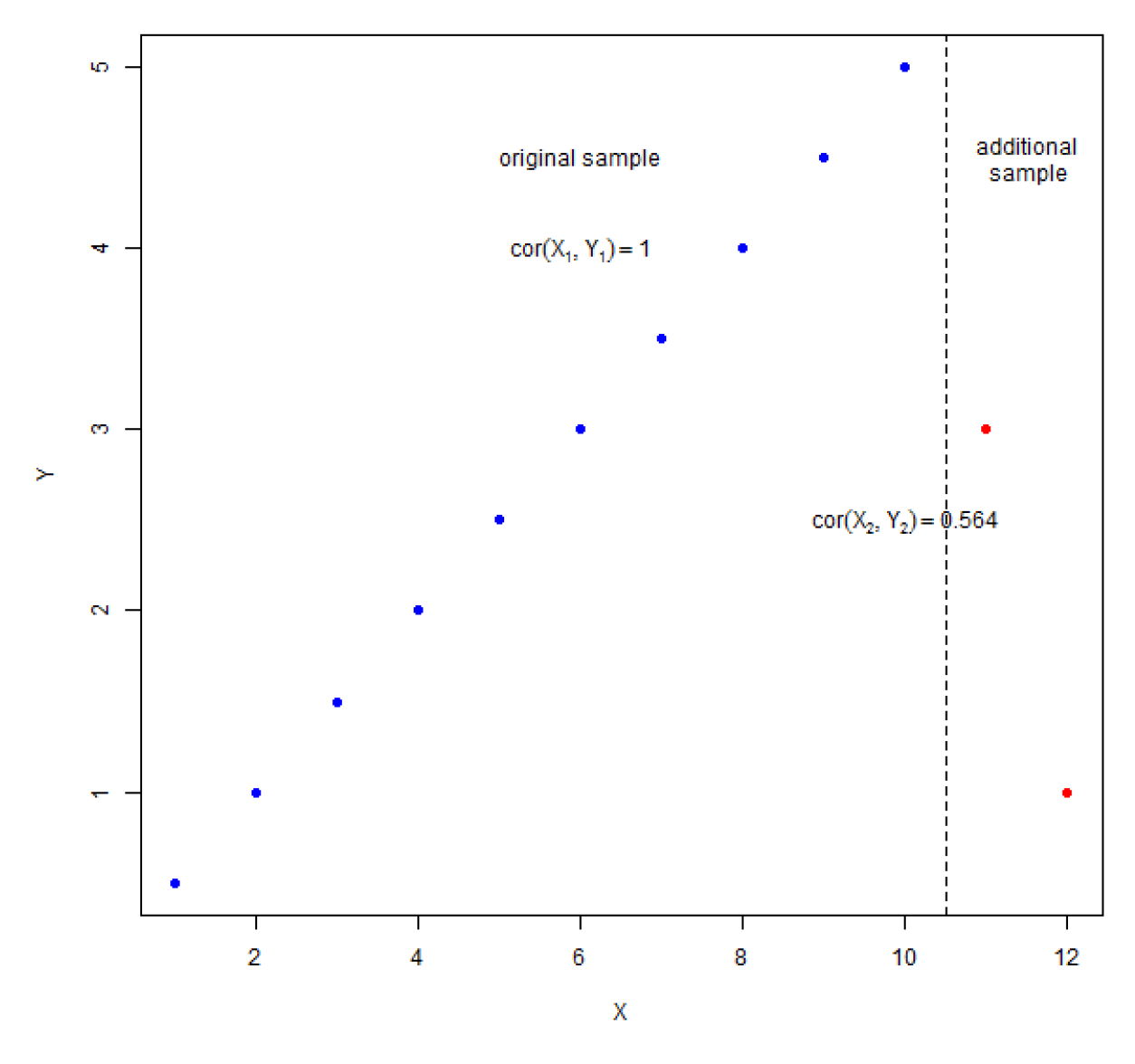
Scatter plot of varibale *X* and *Y*

As Figure 4 shows, we use two threshold values here: *θ*_1_ is the threshold to define *min*(**s**)+ *θ*_1_, less than which is considered as condition specific module. *θ*_2_ is the threshold to define *max*(**s**) − *θ*_2_ greater than which is considered as condition conserved module.

**Figure 4:**
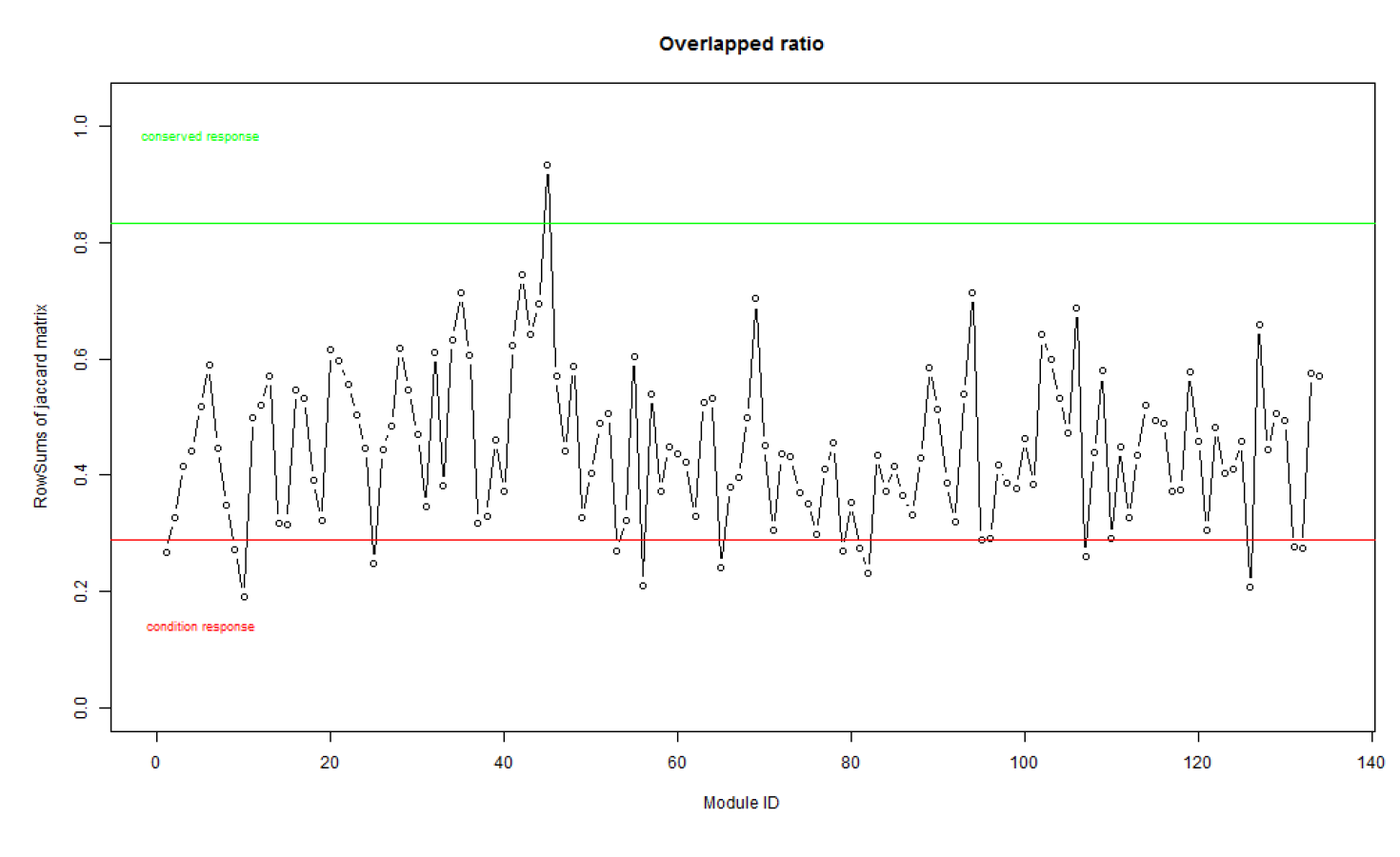
Overlap degree of modules in *N*_1_ with *N*_2_

We also calculate the frequency of each module is annotated as conserved or condition specific and compare all the conditions together. The rationale behind this statistics is based on the mechanism of correlation, i.e. which samples could make an impact on the correlation while others may not? The package visualizes it with a bar plot as Figure 4. A similar plot about the conserved module is also available. The module id is stored as a plain text file for functional enrichment analysis. Here we send one module as gene list to DAVID [10, 11] for integrative analysis.

### 5.2 Evaluation

We evaluate the effectiveness of proposed methods on both synthetic data and real-world data. The basic synthetic gene expression data is generated by the following logic: given desired correlation matrix *C* ∊ ℝ*^n×p^* with *p* genes which has a clear modular structure that all genes are equally divided into 5 groups according to the similarities. Then we conduct the Cholesky decomposition on *C* such that *C* = *LL^T^*, where *L* is the lower triangular matrix. Finally we project *L* on random matrix *A* ∊ ℝ*^n×p^* to get desired gene expression matrix *X* ∊ ℝ*^n×p^*, which has the rough modular structure defined by correlation *C*. Let *n* = 500 and each group has 100 genes in the simulation. In each group, we allocate the gene id from 1−100, 101−200, 201−300, 301−400 and 401−500 respectively. The correlation matrix of genes in *X* is shown in Figure 5. In another matrix *Y*, we merge the last two groups into one by adding more samples to *X*, and the correlation matrix is shown in Figure 7. The we can compare these two networks with proposed method to see which genes were affected. Gene lists in target fold show that modules that contain gene id from 1−100, 101−200 and 201−300 have large overlap with network 2, while module gene id from 301−500 which were merged have least overlap with network 2. The facts are consistent with experimental settings.

**Figure 5:**
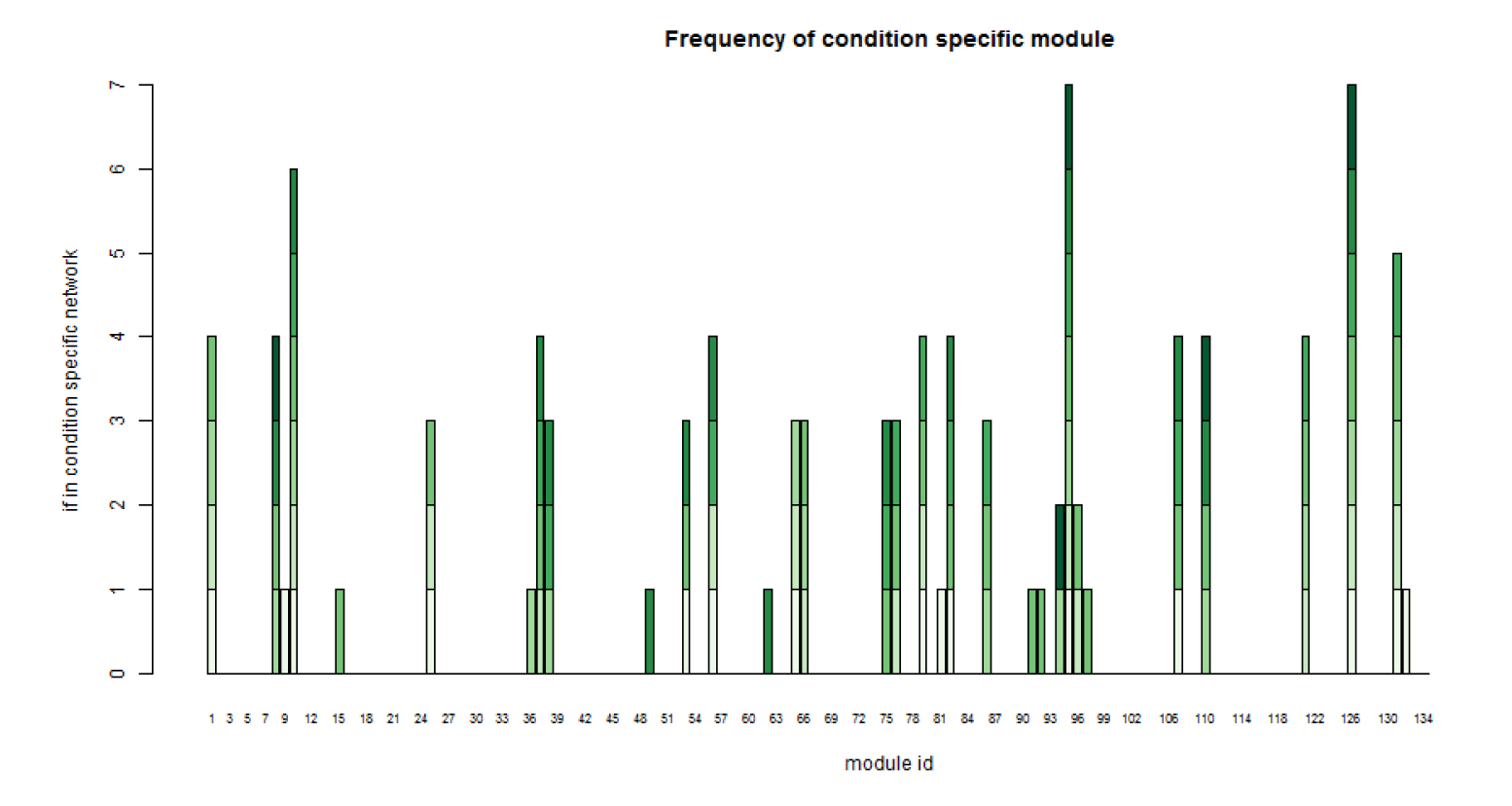
Statistics about which module can be condition specific

Here is the example code to use **MODA** given two gene expression profiles. Results of modules are stored under the newly created folder *ResultFolder* as gene lists. The condition-specific and conserved module ids are stored as plain texts in next directory with the name of indicator which need to be compared. Other materials such as Figure 2 and Figure 4 are also available in the folder.

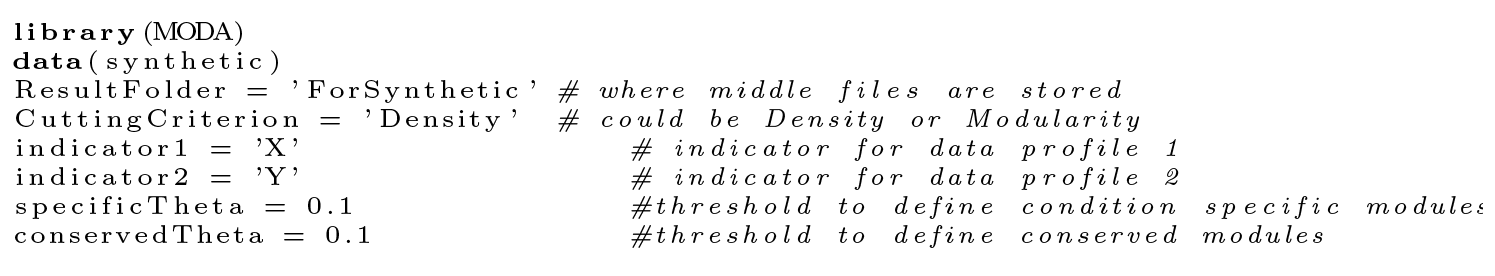

**Figure 6:**
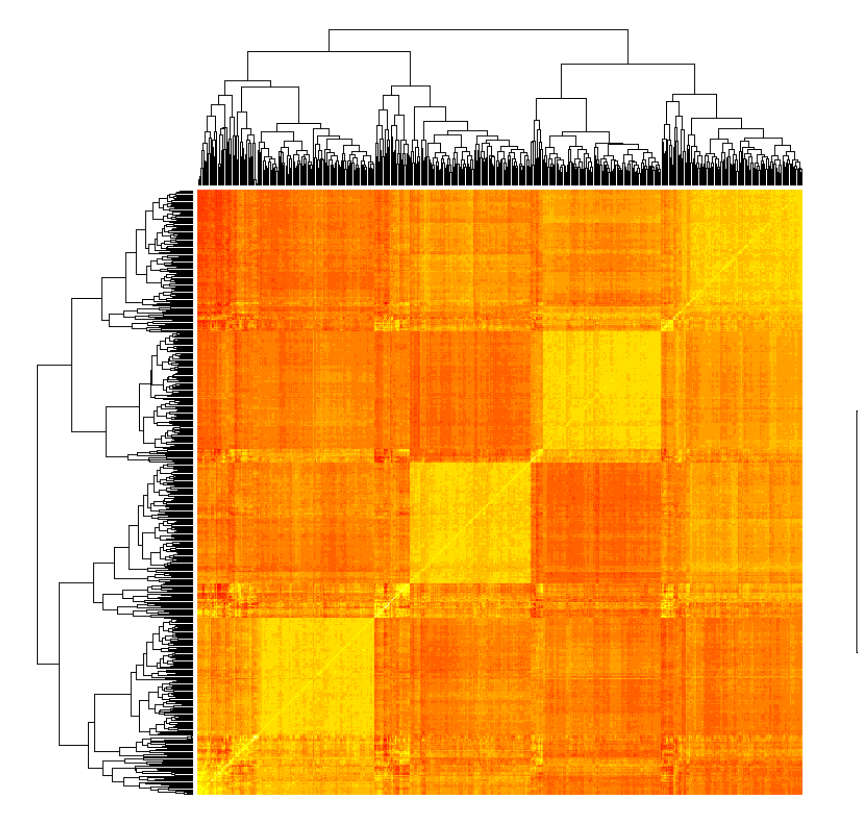
Correlation matrix of *X*

**Figure 7:**
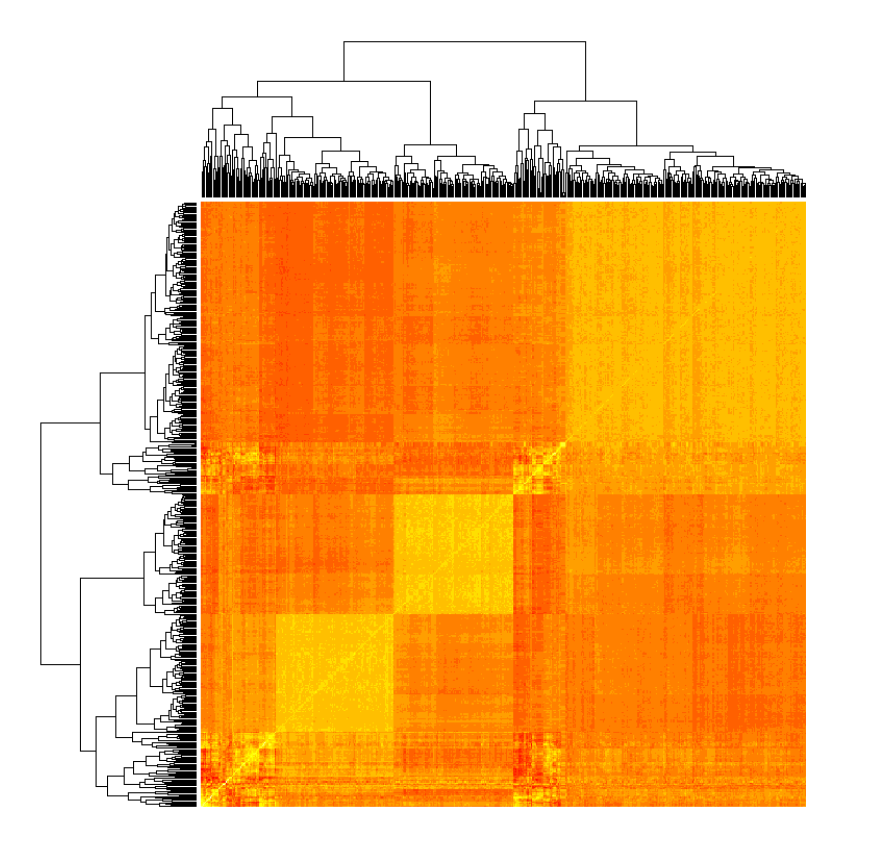
Correlation matrix of *Y*

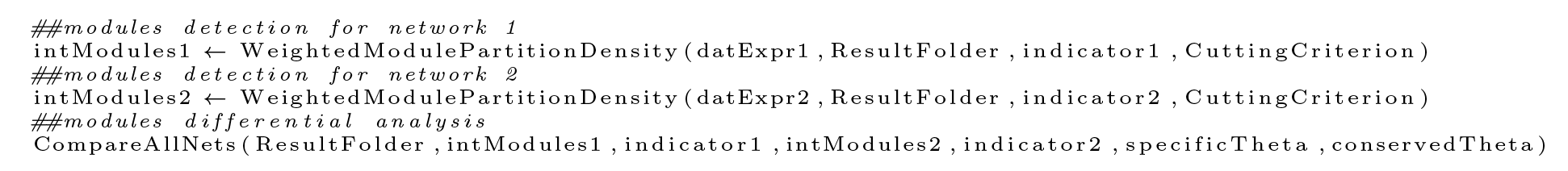

